# FLASH Bragg-peak irradiation with a therapeutic carbon ion beam: first in vivo results

**DOI:** 10.1101/2024.07.12.603197

**Authors:** Walter Tinganelli, Olga Sokol, Anggraeini Puspitasari, Alexander Helm, Palma Simoniello, Christoph Schuy, Sylvie Lerchl, Denise Eckert, Julius Oppermann, Anna Rehm, Stefan Janssen, Denise Engel, Ralf Moeller, Rossana Romano, Claudia Fournier, Marco Durante, Uli Weber

**Affiliations:** GSI Helmholtzzentrum für Schwerionenforschung, Biophysics Department, Darmstadt, Germany; Holland PTC, Delft, the Netherlands; Department of Science and Technology, University of Naples Parthenope, Naples, Italy; Algorithmic Bioinformatics, Justus Liebig University Giessen, Germany; German Aerospace Center, Institute of Aerospace Medicine, Radiation Biology Department, Aerospace Microbiology, Cologne/Köln, Germany; Technische Universität Darmstadt, Institute of Condensed Matter Physics, Darmstadt, Germany; Department of Physics „Ettore Pancini“, University Federico II, Naples, Italy; Technische Hochschule Mittelhessen, Gießen, Germany

**Keywords:** FLASH, UHDR, particle therapy, radiation, carbon ions

## Abstract

**Background and purpose:** In recent years, ultra-high dose rate (UHDR) irradiation has emerged as a promising innovative approach to cancer treatment. Characteristic feature of this regimen, commonly referred to as FLASH effect, demonstrated primarily for electrons, photons or protons, is the improved normal tissue sparing, while the tumor control is similar to the one of the conventional dose-rate (CDR) treatments. The FLASH mechanism is, however, unknown. One major question is whether this effect is maintained when using densely ionizing (high-LET) heavy nuclei.

**Materials and Methods:** Here we report the effects of 20 Gy UHDR heavy ion irradiation in clinically relevant conditions, i.e., at high-LET in the spread-out Bragg peak (SOBP) of a ^12^C beam using an osteosarcoma mouse model.

**Results:** We show that UHDR irradiation was less toxic in the normal tissue compared to CDR while maintaining tumor control. The immune activation was also comparable in UHDR and CDR groups. We observed that the gut microbiome was altered in mice injected with the tumor compared to healthy animals, but both UHDR and CDR exposures steered the metagenome toward a balanced state.

**Conclusions:** The results show that the FLASH effect is safe and effective in heavy ion therapy and provide an important benchmark for the current mechanistic FLASH models.

**Highlights:**

- FLASH irradiation with SOBP carbon ions spares normal tissue in mouse
- Tumor control, immune response, and gut microbioma changes are induced at the same extent both at conventional and ultra-high dose rate
- FLASH carbon ion irradiation is a safe and effective alternative to conventional radiotherapy.

## Introduction

Ultra-high dose rate (UHDR) irradiation has the potential to widen the therapeutic window by reducing the damage to healthy tissues while maintaining the tumor control (commonly referred to as the FLASH effect), as compared to the treatments at conventional dose rates (CDR) (1, 2). The protective effect of FLASH relies not only on the dose rate but also on total dose, spill structure and biological endpoint (1, 3, 4). Several hypotheses proposed to explain the mechanism of the FLASH effect (5). Most models are based on radiation chemistry. The FLASH effect might stem from the rapid generation of free radicals and distinctions in redox and free radical chemistry between normal and tumor tissues, resulting in significantly higher energy deposition and ionization events for a given isodose in UHDR mode compared to CDR mode (6). Another hypothesis suggests that irradiation at UHDR might partially deplete oxygen in the tissues, causing temporal hypoxia an thus increasing their radioresistance (7). Moreover, biological factors might be at play, such as changes in the immune system response to radiation or altered signaling pathways within cells exposed to UHDR (8, 9). UHDR may also differently affect the tumor blood vessels (i.e., causing less damage), potentially preventing changes in tumor vasculature that could affect its response to treatment (e.g., through better immune cells infiltration) (10). Interestingly, all models predicts that the FLASH effect should be LET-dependent, with a tendency to disappear at high-LET (11). Experiments with heavy ions in FLASH conditions are therefore interesting for three reasons: 1. test of the current mechanistic models at high-LET (11); 2. clinical implementation in C-ion therapy (12); and 3. If the sparing effect survives at high-LET, possibility of using very heavy ions (such as ^20^Ne or ^40^Ar) in therapy, and idea originally pursued by Cornelius Tobias at the Lawrence Berkeley Laboratory (13, 14).

We have recently shown the feasibility of UHDR irradiations with carbon ion beams and confirmed the FLASH effect *in vitro* and *in vivo* with heavy ions (15, 16). However, those experiments were performed in the plateau region of ^12^C-ion Bragg curve with relatively low LET values (below 20 keV/μm). In heavy ion therapy, tumors are exposed in the Bragg peak region, which has to be widened to cover the whole tumor volume (Spread-Out-Bragg-Peak; SOBP). In the SOBP, the beam LET is substantially higher (typically 40-80 keV/μm) resulting in an increased biological effectiveness (17), especially for hypoxic tumors (18).

Here, we present the first results of normal and tumor tissue irradiation in the SOBP (20 Gy, LET ≈ 65-85 keV/μm) of ^12^C ions at ultra-high (>100 Gy/s) dose rates accelerated at the SIS18 synchrotron of the GSI Helmholtz Center. The biological model was the LM8 osteosarcoma injected in the hind limb of C3H/He mice. In addition to the measurements of tumor growth and normal muscular tissue damage, we report here on the studies of metastases formation, local and systemic immune responses, as well as changes in gut microbiome.

## Materials and Methods

### Animal model

All experiments were performed using 4-week-old female C3H/He mice (Charles River Laboratories) according to German federal law and with the approval of the Hessen Animal Ethics Committee (Project License DA17/2000). A total of forty-four mice were used for the experiments. Mice were divided into four groups: UHDR (14 animals), CDR (15 animals), sham irradiated (10 animals), and no tumor (5 animals). The mice were housed at GSI in a conventional animal facility (non-SPF), at ∼22°C and a 12-hour light-dark cycle. They were provided with unrestricted access to a standard diet (Ssniff, Germany) and water. Seven days before irradiation, mouse Dunn osteosarcoma LM8 cells (originating from C3H/He mice, purchased from Riken BioResource Center (Japan)) were injected subcutaneously in the right posterior limb of the mice (10^6^ in 20 µl of DMEM medium with 10% FBS and 1% Pen/Strep). After one week, the tumors were palpable and measurable.

### Irradiation

Irradiations were performed in the Cave A branch of the SIS18 synchrotron at the GSI Helmholtz Center in Darmstadt, Germany. A beam of 240 MeV/u ^12^C-ions was extracted in air. The beam line and setup were similar as shown and described in (19), while the general concept of the dosimetry system in the vault is described in (20). However, for the UHDR irradiation some modifications were applied (see supplementary materials).

Mice were anesthetized with isoflurane and then irradiated (20 Gy) inside an airtight plastic box flushed with isoflurane at a specific rate to maintain anaesthesia. Since the irradiation field, particularly the lateral dose fall-off, exceeded the target volume, we shielded the mice bodies with brass absorbers to prevent side effects in adjacent organs (Figure 1A). The positions of the limbs were adjusted 10 mm before the distal edge of the SOBP using a plastic absorber of 60 mm water equivalent thickness. Due to the strong gradient, the dose-averaged LET in the target volume varied between 65-85 keV/μm (Figure 1B).

**Figure 1.**
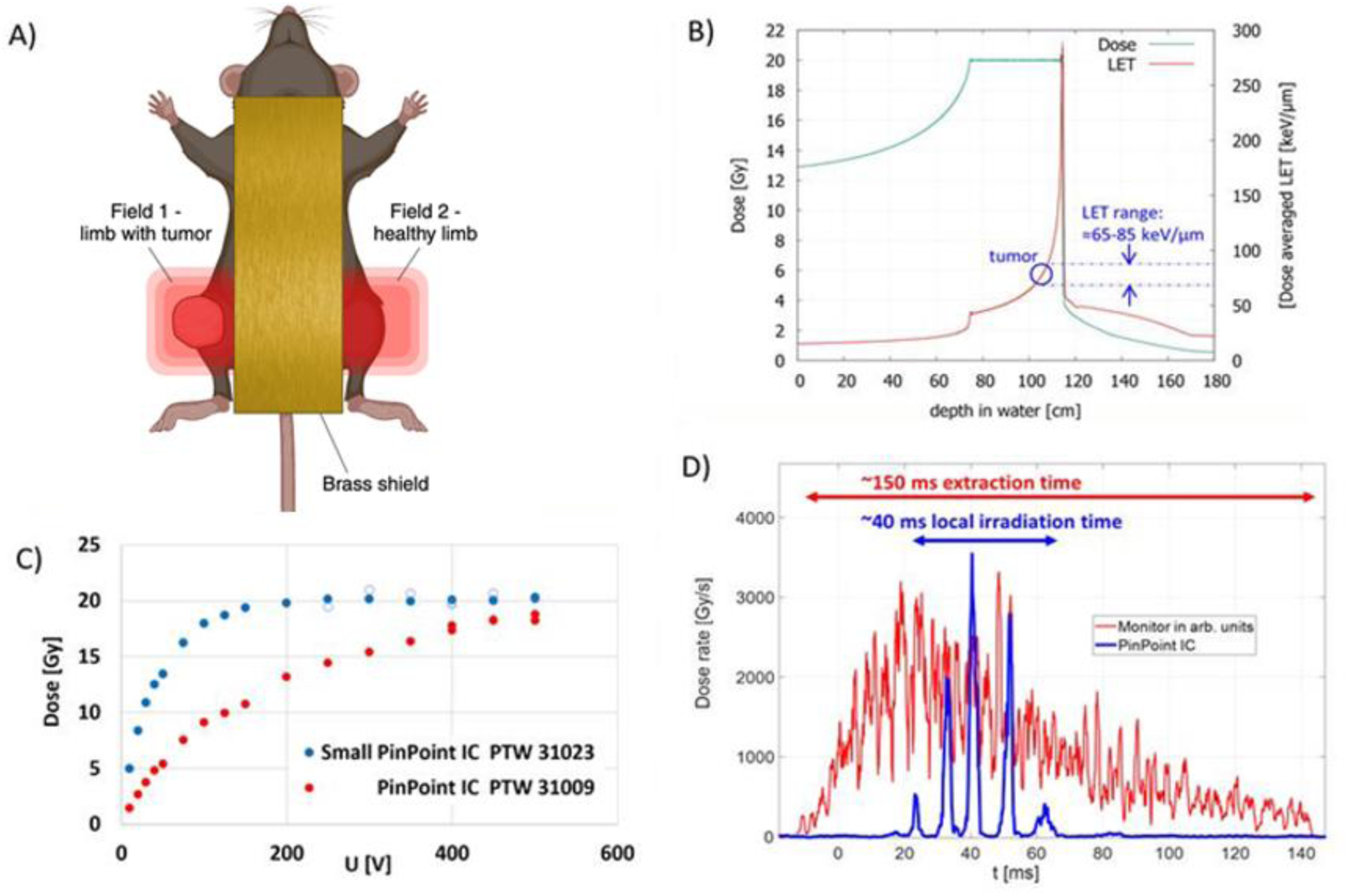
Setup for the FLASH (UHDR) and CDR irradiation. A: Mice irradiated successively with two 20 Gy fields (either CDR or UHDR, ≈15 × 15mm^2^ homogeneous area) for the tumour-bearing limb and the healthy limb. Adjacent organs were shielded by a 25 × 80 mm^2^ and 10 mm thick block of brass. B: Depth-dose and LET distribution from the 240 MeV/u ^12^C beam modulated by the 2D-RM into a 4 cm SOBP. Tumour depth is adjusted at 10 mm before distal edge by absorbers. C: Saturation curves at UHDR conditions for two different PinPoint^TM^ chambers, type 31023 chamber was selected for dosimetry. D: Time resolved signal and local dose rate (mid of target volume) from the beam monitor and the PinPoint IC (PTW 31023), respectively, showing the total and local irradiation time at the center of the target.

The treatment field was applied by the Cave A raster scanning system with a scanning pattern of 21×21 mm^2^ (8×8 beam spots, step size of 3 mm) and a pencil beam of ≈6mm FWHM, yielding a highly homogenous dose in the inner area of 15×15 mm^2^. The SOBP with an extension of 4 cm was realized by a so-called 2D range modulator (2DRM) (19) with a pin period of 2 mm and a pin height of ca. 40 mm. It was placed in a distance of 50 cm from the target in order to avoid the edge scattering inhomogeneities from the modulator (21). The modulator was precisely printed with a stereo lithography (SLA) 3D-printer (3D-Systems). Additionally, a standard 3 mm ripple filter (22) was installed at the exit window to further smoothen the depth dose profile.

### Tumor growth measurements

Tumor size was measured with a Vernier caliper approximately every second day for 24 days after irradiation. Mice were sacrificed 35 days after tumor inoculation. Tumors had approximately hemi-ellipsoid shapes and therefore their volumes were calculated from the measured length (a), width (b), and depth (c) as V = (π × a × b × c) / 6. Statistical analysis was done using a T-test between the different groups.

### Histological analysis

Twenty-eight days after irradiation, animals were sacrificed and the primary tumor, lungs, spleen and both limbs were collected for further analysis. Limb muscles and tumors were fixed in 4% neutral buffered formalin for 24 hours. After 24 hours, the specimens were dehydrated in a Tissue Processor (Leica HistoCore Pegasus) according to the manufacturer settings, embedded in paraffin (Leica; HistoCore Arcadia Embedding Center) and sequentially cut and stained with hematoxylin (Gill No.3, Sigma Aldrich) and 0.5% aqueous eosin solution (Carl Roth) according to the manufacturer protocol. To assess the level of fibrosis in limb muscles, Picrosirius Red staining according to (23) was performed. Using a microtome (RM2235, Leica Microsystems), tumors were cut into 5 µm slices. To stain for CD8 alpha positive cells, tumor slices were processed with rabbit recombinant anti-CD8 alpha (monoclonal, ab209775, Abcam) as primary antibody as previously described (24, 25). Subsequently, a digitizer for slides (Axio Scan.Z1, Carl Zeiss Microscopy) was used to image 2-3 entire slices of each tumor (n = 5-11 tumors per treatment group). Further details are given in the Supplementary material.

### Lung metastasis analysis

After the animals were sacrificed, the bilateral lungs were collected and placed in Bouin solution (Sigma-Aldrich) for 24 hours. An operator with a stereoscope counted the metastatic nodules on the pulmonary surfaces. The average incidence of lung metastases was expressed as relative values compared with that of the control group (25). Statistical analysis was performed via Mann-Whitney test using the GraphPad Prism 9 software.

### Flow cytometry analysis

To obtain single cell suspensions of spleen and lymph nodes, the isolated organs were manually disrupted using tissue strainers (40 or 100 µm; Greiner Bio-One). A subsequent lysis of red blood cells was performed for 5 minutes in a buffer containing ammonium chloride, potassium hydrogencarbonate and EDTA disodium salt. Afterwards, cell suspension was washed in PBS, centrifuged, and resuspended in a medium consisting of RPMI 1640, FCS, MENEAA, Pen/Strep, L-Glutamin and sodium pyruvate. The number of viable cells per ml were counted by using 0.4% trypan blue solution (ThermoFisher Scientific) and an automated cell counter (TC20™, Bio-Rad Laboratories). 2 × 10^6^ cells per sample were separated for each staining panel. After blocking the cells in a buffer containing mouse serum, FCS and optional Fc-Block (BD Bioscience), cell surface proteins of interest were labeled with respective antibodies according to manufacturer’s instructions. Optionally, cells were subsequently fixed and permeabilized for additional intracellular staining as specified by the manufacturer. The immune cell distribution was measured with a CytoFLEX S flow cytometer (Beckman Coulter) and data were analyzed with the software Cytexpert v2.4 (Beckman Coulter). Panels and analysis are described in the Supplementary material.

### Microbiome – sampling, DNA isolation and sequencing

At the experimental endpoint, gut tissues from all the mice were extracted, fixed and stored at - 80°C in DNA/RNA Shield solution (Zymo Research) according to manufacturer’s instructions. The samples were processed and analyzed with the Microbiome Analysis Service: Targeted Metagenomic Sequencing (Zymo Research Europe, Freiburg, Germany). Details are provided in the Supplementary materials.

### Bioinformatic analyses – gut microbiome

Demultiplexed paired end fastQ reads have been directly downloaded from Zymo Research. Primer sequences (fwd=CCTAYGGGDBGCWGCAG, rev=GACTACNVGGGTMTCTAATCC) have been trimmed off the raw reads via cutadapt (v4.2) (26). Following processing only used forward reads (R1) which were further trimmed to exactly 150bp from which we called Amplicon Sequencing Variantes (ASVs) via Deblur (v1.1.1) (27) through the Qiita platform (28). ASVs with less than 10 reads in all samples together were dropped. ASVs have been phylogenetically placed into the Greengenes 13.8 99% backbone tree via SEPP (v4.3.10). The resulting tree was used to compute weighted and unweighted UniFrac (29) and Faith’s PD alpha diversity (30). Taxonomy was assigned via Qiime2’s “feature-classifier” plugin version 2021.4 against Greengenes 13.8 99% OTU sequences (31). Bacterial features with assigned taxonomy containing the labels c Chloroplast or f mitochondria were considered of host or plant origin and removed from the feature table prior to rarefaction. We rarefied the ASV table to 35000 reads per samples without losing any. The final ASV table comprised 530 ASVs and 42 samples.

## Results

In our irradiation setup we exposed both limbs in the SOBP of the carbon ion beam (Figure 1). We could therefore access the tissues from both the healthy limb and the one where the tumor was implanted.

### UHDR ^12^C irradiation reduces damage to healthy tissues

To assess the impact of UHDR and CDR ^12^C exposure on healthy tissues inside the SOBP, we investigated fibrosis formation by collagen deposition using Picrosirius Red staining of the irradiated muscle tissues (23, 32). Collagen deposition was significantly lower following the UHDR irradiation. Notably, the distribution and the localization of Picrosirius Red staining in the FLASH-irradiated muscles closely resembled that of unirradiated muscle tissue, as shown in Figure 2A. The following quantification demonstrated a significant level of collagen I deposition following CDR ^12^C irradiation compared to UHDR irradiation, as shown in Figure 2B.

**Figure 2.**
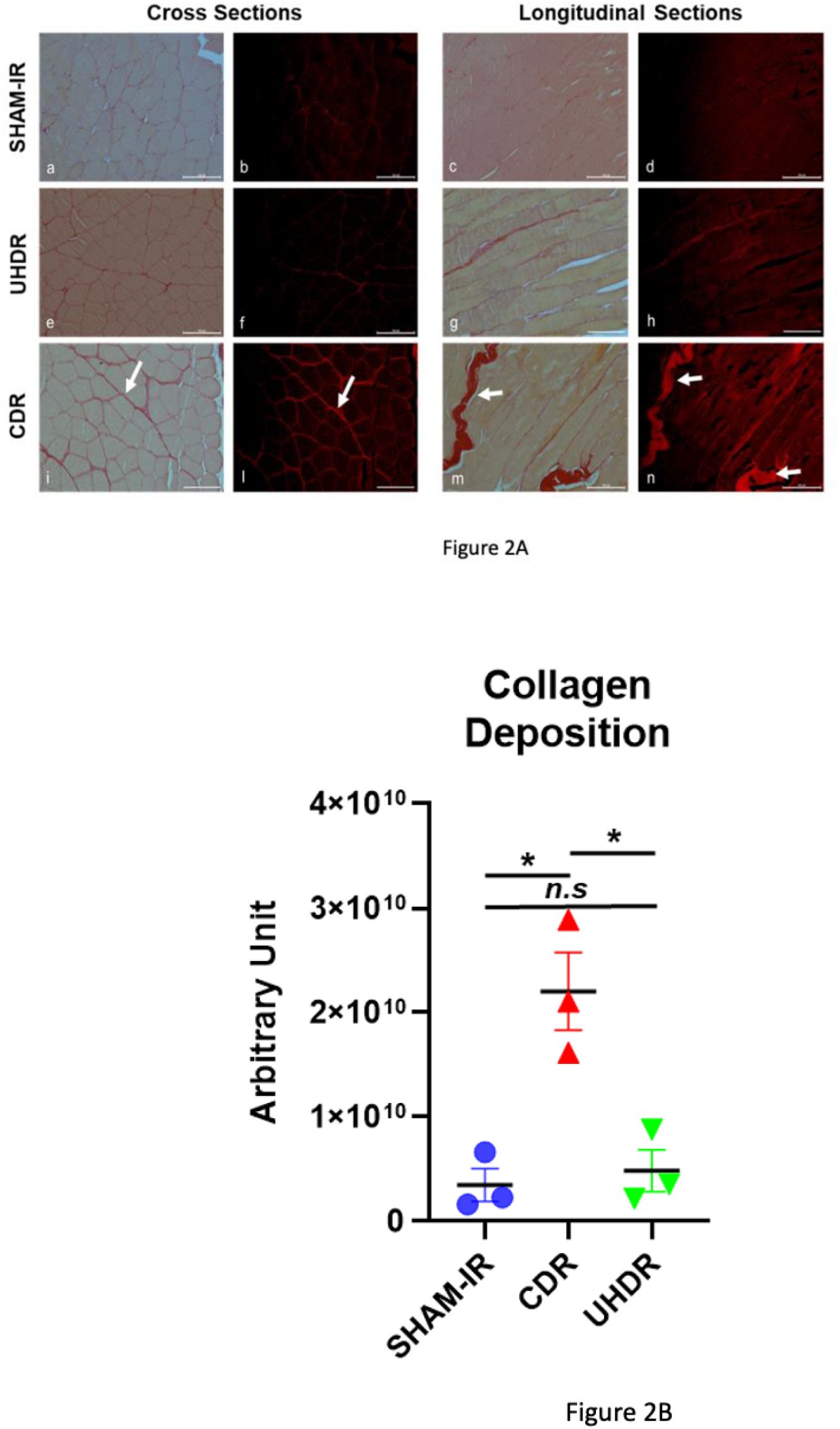
Muscular damage inside the irradiated healthy limb following UHDR and CDR irradiation. A: Fibrosis and collagen I deposition were observed in longitudinal and cross sections of healthy muscle tissue stained with Picrosirius red in sham control (a-d), UHDR-irradiated (e-h), and CDR-irradiated (i-n) mice. The distribution and localization of collagen I in the UHDR-irradiated samples were similar to those in the control mice. A high amount of collagen I was observed between the myofibers (indicated by white arrows) following CDR irradiation. Scale bars: 100 μm. B: Quantification of collagen I deposition. Blue circles represent sham-irradiated mice (n=9), red triangles represent CDR-irradiated mice (n=9), and green triangles represent UHDR-irradiated mice (n=9). The bars represent the mean values (± standard error of the mean, SE). *n.s:* not significant; * P < 0.05.

### UHDR and CDR ^12^C irradiations lead to equivalent tumor control

The average tumor volume as a function of post-irradiation time is illustrated in Figure 3A. Both UHDR or CDR exposure at 20 Gy led to equivalent tumor control, while progressive tumor growth is observed in sham irradiated animals.

**Figure 3.**
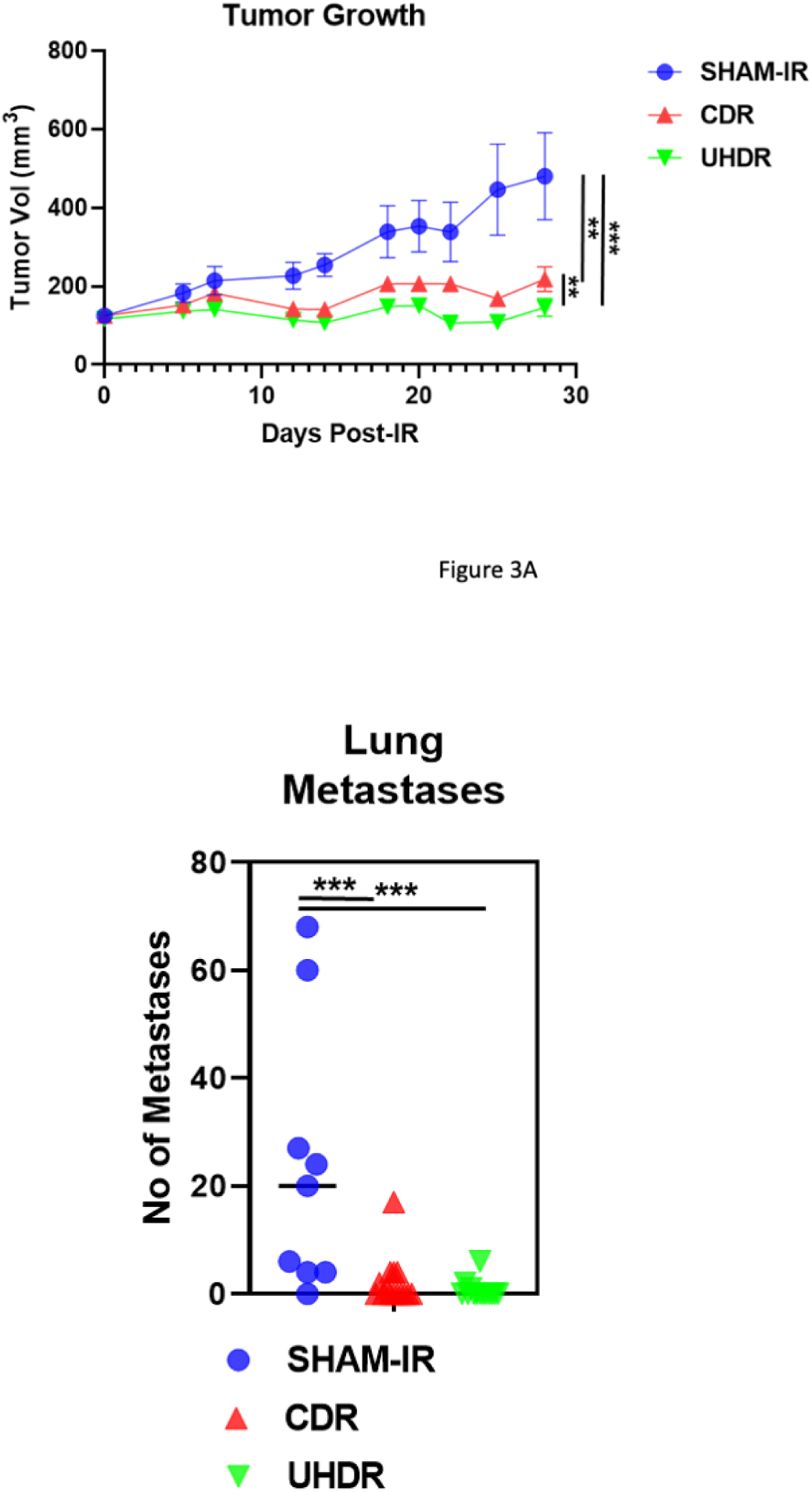
Comparative Analysis of Tumor Volume Reduction and Metastatic Nodule Reduction Following CRD and UHDR Irradiation. (A) The reduction of tumor volume is presented over a span of 28 days post-irradiation. Blue circles represent sham-irradiated mice (n=10), red triangles represent CDR-irradiated mice (n=15), and green triangles represent UHDR-irradiated mice (n=14). Black bars indicate mean values ± SEM for each treatment group. (B) The reduction in the number of metastatic nodules following CDR and UHDR irradiation is shown. Similar to Figure 2, blue circles represent sham-irradiated mice (n=10), red triangles represent CDR-irradiated mice (n=15), and green triangles represent UHDR-irradiated mice (n=14). Black bars also indicate mean values ± SEM for each treatment group. ** P< 0.005; *** P< 0.001.

### UHDR and CDR ^12^C irradiations lead to suppression of lung metastases

The LM8 osteosarcoma tumor in the hind limb strongly metastasize in the lung of the mice. We observed a significant decrease in the number of lung metastases in both treatment groups (UHDR: 0.6 ± 0.4; CRD: 2 ± 1) compared to the control group (24 ± 8) (Figure 3B). We thus confirmed the robust efficacy of CDR in suppressing lung metastases reported in our previous study (25), and we believe that it can explain the lack of significant differences between the UHDR and CDR modes observed in the present work.

### UHDR ^12^C irradiation does not differentially impact on local and systemic anti-tumor immune responses 28 days following exposure

To investigate the effects of UHDR ^12^C irradiation on the local immune response in the tumors, we analyzed the number of cells stained positive for CD8, a marker for cytotoxic T-cells, 28 days following exposure. Both UHDR and CDR ^12^C led to a significant increase in CD8-positive cells compared to the controls (Figure 4A). This indicates increased infiltration of cytotoxic T-cells upon irradiation. However, no significant differences in the number of CD8-positive cells were found between UHDR and CDR, suggesting no increased efficiency of UHDR ^12^C with respect to a local radiation-induced anti-tumor immune response at 28 days after the treatment.

**Figure 4.**
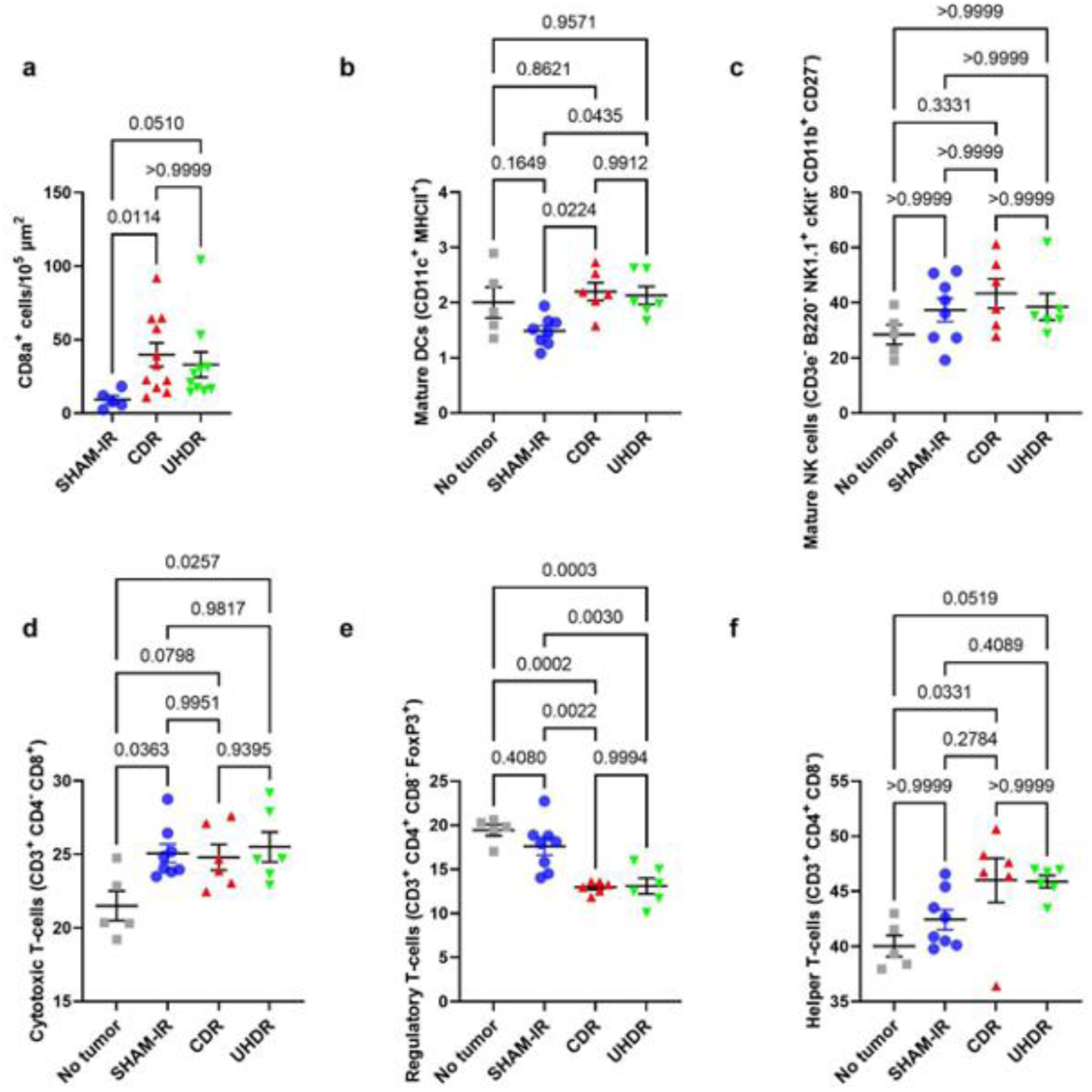
Infiltration of CD8a^+^ cells in tumors and splenic immune cell populations 28 days following CDR or UHDR treatment. A. Numbers of CD8a^+^ cells were normalized to an area of 10^5^ µm^2^ for each analyzed tumor; black bars indicate mean values ± SEM for each treatment group; SHAM-IR n=5, CDR n=11, UHDR n=10. B-F) Percentages of splenic immune cell populations; No tumor n=5, SHAM-IR n=8, CDR n=6, UHDR n=6. Black bars indicate mean values ± SEM for each treatment group for B: mature dendritic cells, C: mature NK cells, D: cytotoxic T-cells, E: regulatory T-cells and F: helper T-cells. The p-values (Kruskal-Wallis test) are indicated individually.

Next, to examine whether UHDR and CDR differentially impact the potential to induce systemic immune responses with putative effects on metastases, we analyzed immune cell populations in the spleen and lymph nodes. We screened for dendritic cells (DCs), B-cells, NK cells in spleen and several subpopulations of T-cells in spleen and lymph nodes. Analyses of the main populations of immune cells in spleen and lymph nodes revealed no significant changes (Figures S1 and Figure S2A). By tendency, however, sham-irradiated tumor-bearing mice showed slight differences in the percentage of immune cells compared to no-tumor mice, which were partly reversed by both irradiation modes. The only significant radiation-induced increase was found in splenic NK cells following CDR exposure (Figures S1C).

The amount of splenic mature DCs (CD11c+ MHCII+ cells, Figure 4B) was found to be significantly (p<0.05) increased after exposure to both radiation modes, CDR and UHDR, compared to sham-irradiated tumor-bearing mice, but no differences between the modes occurred. In contrast, the percentage of mature NK cells (CD3-CD19-CD11b+ CD27-cKit-NK1.1+ cells, Figure 4C) did not reveal any significant differences but, again, showed a tendency to increase in tumor-bearing mice (both irradiated and sham-irradiated). This was the same for cytotoxic T-cells (CD3+ CD8+ CD4-) but the change differed significantly from no-tumor mice (Figure 4D). Radiation-induced changes, both compared to healthy and sham-irradiated mice, were found for regulatory T-cells (CD3+ CD4+ CD8-FoxP3+) and helper T-cells (CD3+ CD8-CD4+) (Figures 4E and 4F). While the number of regulatory T-cells significantly decreased compared to both controls (Figure 4E), the amount of helper T-cells increased, however, significantly only when CDR was compared to no-tumor mice (Figure 4F). Yet again, no significant differences between the two modes, CDR and UHDR, were found. When the respective T-cell subpopulations were analyzed in lymph nodes, no differences were found for cytotoxic T-cells and helper T-cells (Figures S1B, C) but the results for regulatory T-cells stemming from spleen were confirmed by tendency (Figure S1D).

### UHDR and CDR ^12^C irradiations push the microbiome back towards the healthy state

Taxonomic composition of the gut microbiome (Figure 5A) and pairwise beta diversity distances (Figure 5B) was assessed via 16S rRNA profiling. We found that the presence of limb osteosarcoma modifies the gut microbiome in mice, as indicated by statistically significant differences between the no-tumor mice and tumor-bearing, sham-irradiated mice (Figure 5C). In fact, these eight untreated mice had a significantly different gut microbiome from the irradiated tumor-bearing mice (Figure 5D). It remains unclear whether this change is caused directly by off-target irradiation of the gut and subsequently gut bacteria, or indirectly by more excessive outgrow of tumor burden in the untreated mice, or a combination of both. Nevertheless, the gut microbiome of the irradiated mice resembled that of the healthy mice significantly closer than the microbiome of the sham irradiated animals for both treatment regimens (Figure 5E, p<10^-7^). We could furthermore detect significant difference between UHDR and CDR treatments (Figure 5F); however, it did not lead to significantly closer microbiomes compared to the healthy mice. In conclusion, presence of tumors causes or reflects a presumably detrimental microbial shift, which can partially be reversed by irradiation treatment. UHDR and CDR irradiation modes impact the microbiome differently, however both push the microbiome back towards the healthy state, with indiscernible intensity.

**Figure 5.**
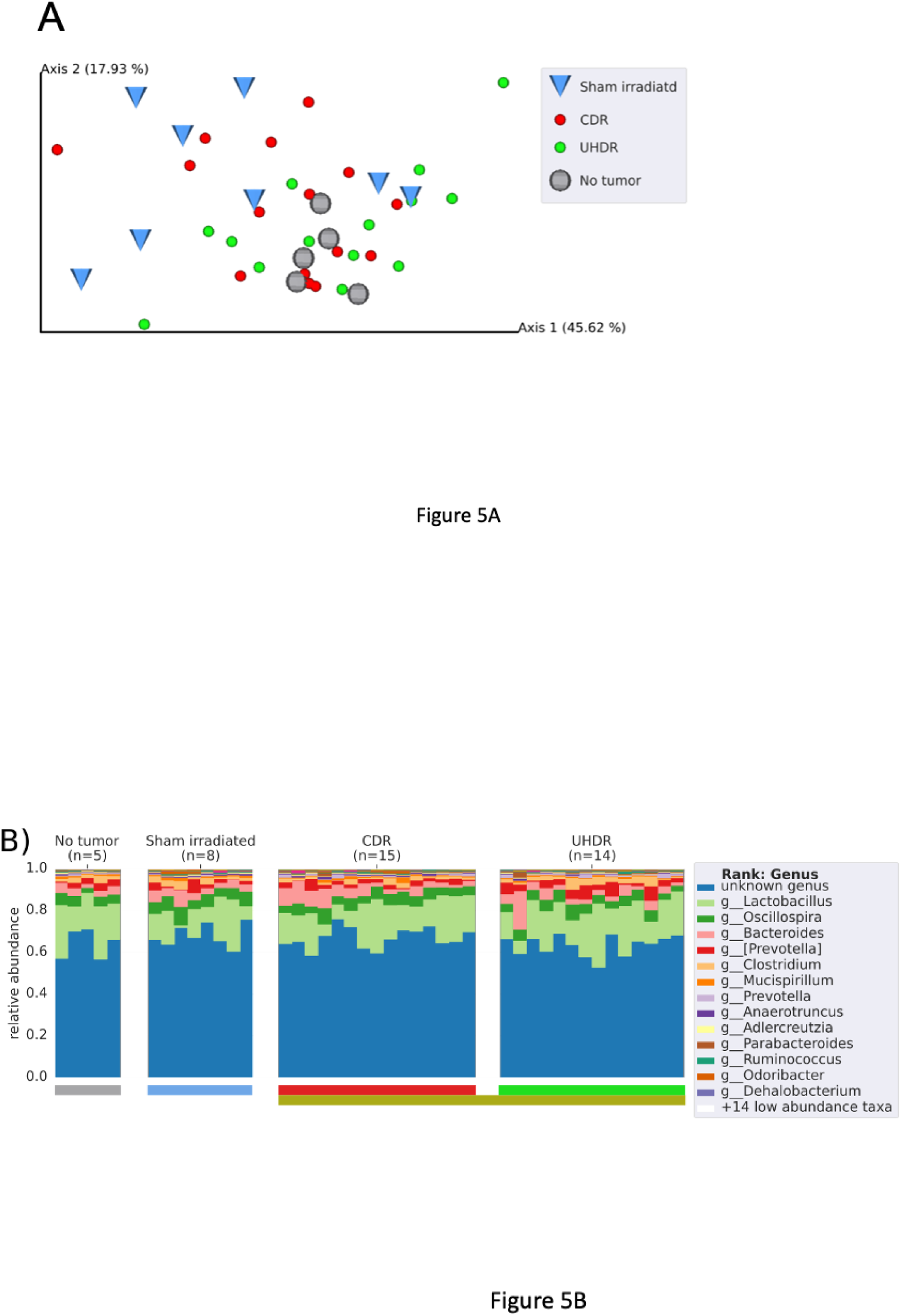
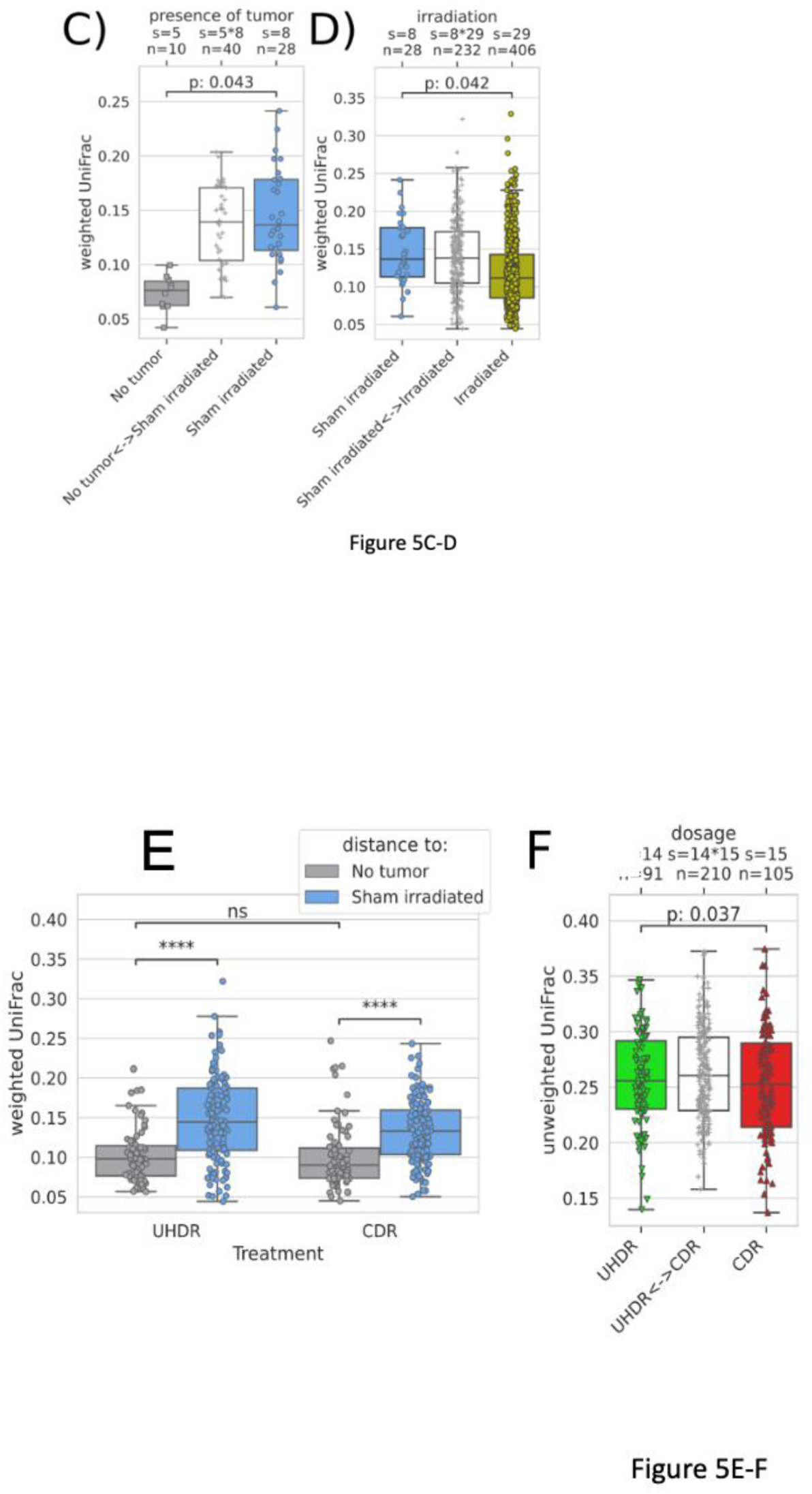
Changes in the gut microbiome caused by the presence of tumor and by different irradiation modes. A: PCoA ordination of pairwise un-weighted UniFrac distances. B: Taxonomy bar plot on genus rank. C-D, F: visualizations of PERMANOVA tests which consider differences within group (left and right box) and across group (white box in the middle). E: distances from UHDR, CDR to control, (no tumor) group (gray) and to sham irradiated tumor-bearing mice (blue). Distances are compositional, i.e., low distance to gray implies high distance to sham and vice versa.

## Discussion

We report here the first experiment on the FLASH effect in an animal model irradiated in the SOBP of a heavy ions beam. We observed that animals irradiated at CDR within the ^12^C SOBP had increased collagen deposition, a hallmark of radiation-induced fibrosis in the healthy muscle tissues. In contrast, those irradiated at UHDR had collagen levels similar to those in non-irradiated control animals. This confirms the sparing effect of UHDR radiation, even with high-LET ^12^C beams. This result is valued for testing the current mechanistic models of the FLASH effect, which generally predict a decrease of the effect at high-LET (11).

Similar to our previous low-LET C-ions (16), we found no significant differences between the CDR and UHDR groups in tumor control. The measurements of tumor sizes showed that the UHDR-irradiated tumors were slightly smaller than those irradiated at CDR. This size reduction could potentially be attributed to the inflammation in the healthy tissues surrounding the tumor, caused by CRD but not UHDR irradiation.

In our previous work (16) we observed a decrease in the occurrence of lung metastases following ^12^C UHDR irradiation. In the SOBP, both CDR and UHDR exhibited equal efficacy in suppressing the lung metastasis formation. The ability of high-LET ^12^C ions to significantly inhibit metastatic development at CDR has already been demonstrated in our earlier study (25). Looking at the generalized immune response, both UHDR and CDR irradiations led to a significant increase of infiltrated CD8+ cytotoxic T-cells in the tumour and a decrease of Treg cells, but no significant differences were observed between CDR and FLASH. Taken together, these data suggest that SOBP (high-LET) C-ions elicit a strong immune response, and UHDR conditions do not modify it. The impact of different radiation strategies on the microbiome is an area of ongoing research and investigation. Some preliminary research has suggested that radiation therapy, especially when administered at high doses (which are indeed a requirement for the FLASH effect), may lead to alterations in the gut microbiome (33).

To assess potentially detrimental effects of the irradiation on the gut microbiome of mice, we profiled their bacterial colon content via 16S rRNA amplicon sequencing. Our results suggest that tumors negatively affect the gut microbiome. However, when treated with irradiation, regardless of the dose rate, this shift can be partially reversed. Both treatments tend to guide the microbiome back to a healthier state.

In conclusion, our results show that normal tissue sparing and tumor control are maintained when high-LET heavy ions are used at UHDR in a murine osteosarcoma model. On the other hand, the immune response and alterations in the gut microbiome were induced by SOBP C-ions but were not dependent on dose rate. These results are relevant for understanding the mechanisms of the FLASH effect, since all the present models are ostensibly affected by the radiation LET. Along with our previous results in plateau-phase C-ions (16), they demonstrate the feasibility of UHDR C-ion radiotherapy.

## Acknowledgments

This work was funded by the European Union’s Horizon Europe research and Innovation programme under Grant Agreement No. 101057511 (EURO-LABS).

## Statistical analyses

AR, SJ (bioinformatics), MD, OS (other data)

## Supplementary materials and figures

### Irradiation setup and dosimetry

To achieve FLASH conditions, the total dose of 20 Gy (±0.3 Gy fluctuations in σ) was delivered in a single synchrotron spill of ca. 150 ± 20 ms containing 5-7×10^9^ ^12^C ions. Under these conditions, the average dose rate calculated as target dose per total irradiation time reached >100 Gy/s, while the local dose rate due to the raster scanning irradiation was up to >3000 Gy/s (instantaneous dose rate) and >300 Gy/s for the threshold (2.5-97.5%) related local dose rate (Figure S3D and ref. (11)). To enable a precise dose delivery and to avoid saturation effects, the parallel plate ionisation chamber for beam monitoring was operated with a He-Argon (80/20%) mixture (34) and the dose was measured by a mini-Farmer chamber (PinPoint™ type PTW 31023) operated at 400 V. Due to a high field strength and a small active volume this chamber does not show saturation effects at UHDR (Figure S3C). For CDR irradiation, intensity was lowered and animals were exposed to multiple spills at a dose rate around ∼20 Gy/min in a typical time of 60 s.

### Histological analysis

To quantify the amount of collagen I in the healthy muscle of the animals, we quantified the intensity of the red florescence signal (Nikon Eclipse Ni-U microscope, camera Nikon DS-Fi3, NIS-elements F 5.21.00 64-bit software). A total of nine animals were included in the study, with three animals assigned to each of the groups (UHDR, CDR and sham irradiated). For each animal, we captured five fields or pictures of longitudinal section. Using ImageJ software, we measured the intensity of the red signal in each field and calculated the average intensity and standard error of the mean for each sample. Statistical analysis was done using a T-test between the different groups.

To quantify the CD8 alpha positive cells, the QuPath software (v0.4.3, GitHub Inc) was used. In each scan, 3-4 random but representative sections of a certain area (approx. 3 × 10^5^ µm^2^) were chosen to count all CD8 alpha positive cells, and the amount was normalized to 10^5^ µm^2^ of tumor cross section. Statistical analysis was performed via Kruskal-Wallis test since normality tests showed no Gaussian distribution of all datasets.

### Flow cytometry

DC panel: anti-CD11c (BV510, HL3, BD Biosciences, 562949), anti-MHCII (APC, M5/114.15.2, BioLegend, 107614); T cell panel: anti-CD3e (PE-Cy7, 145-2C11, BD Pharmingen, 552774), anti-CD4 (Pacific Blue, RM4-5, BD Pharmingen, 558107), anti-CD8 (PE, 53-6.7, BD Biosciences, 553032), anti-FoxP3 (VioB515, REA788, Miltenenyl Biotec, 130-111-603); NK panel: anti-CD3e (PerCP, 145-2C11, Biolegend, 100326), anti-B220 (Pacific Blue, RA3-6B2, BD Pharmingen, 558108), anti-NK1.1 (PE-Cy7, PK136, BD Pharmingen, 552878), anti-cKit (APC, 2B8, eBioscience, 17-1171-83), anti-CD11b (FITC, M1/70.15.11.5, Miltenyi Biotec, 130-081-201), anti-CD27 (PE, LG.3A10, BD Pharmingen, 558754); B Cell panel: anti-CD19 (FITC, MB19-1, eBioscience, 11-0191-82), anti-B220 (Pacific Blue, RA3-6B2, BD Pharmingen, 558108).

Statistical analyses were performed via one-way Anova when normality test indicated Gaussian distribution of all datasets or via Kruskal-Wallis test when no Gaussian distribution for at least one dataset was shown.

### Microbiome

DNA extraction was done using the ZymoBIOMICS®-96 MagBead DNA Kit (Zymo Research, Irvine, CA). The DNA samples were prepared for targeted sequencing with the Quick-16S™ NGS Library Prep Kit (Zymo Research, Irvine, CA). The Quick-16S™ Primer Set V3-V4 (Zymo Research Europe, Freiburg, Germany) was chosen based the best coverage of the 16S gene while maintaining high sensitivity. The primer sets used in this project are marked below. PCR reactions were performed in real-time PCR machines to control cycles and limit PCR chimera formation. The final PCR products were quantified with qPCR fluorescence readings and pooled together based on equal molarity. The final pooled library was cleaned up with the Select-a-Size DNA Clean & Concentrator™ (Zymo Research, Irvine, CA) and quantified with TapeStation® (Agilent Technologies, Santa Clara, CA) and Qubit® (Thermo Fisher Scientific, Waltham, WA). The ZymoBIOMICS® Microbial Community Standard (Zymo Research, Irvine, CA) was used as a positive control for each DNA extraction, if performed, while negative controls (i.e., blank extraction control, blank library preparation control) were included to assess the level of bioburden carried by the wet-lab process. The library was sequenced on Illumina® MiSeq™ with a v3 reagent kit (600 cycles) and 10% PhiX spike-in.

Sequencing data is available either as the public study 15299 in qiita.ucsd.edu or through ENA access. Microbial analysis notebook is available at: https://github.com/jlab/microbiome_Tinganelli_flash.

The following statistical analyses were performed to evaluate the differences between experimental groups: PERMANOVA with 999 permutations of weighted UniFrac distances (Figures 4C,D), two-sided Mann-Whitney on weighted UniFrac (Figure 4E), PERMANOVA with 999 permutations of unweighted UniFrac distances and two-sided Mann-Whitney on Bray-Curtis, weighted and unweighted UniFrac distances (Figure 4F).

---

**Figure S1.**
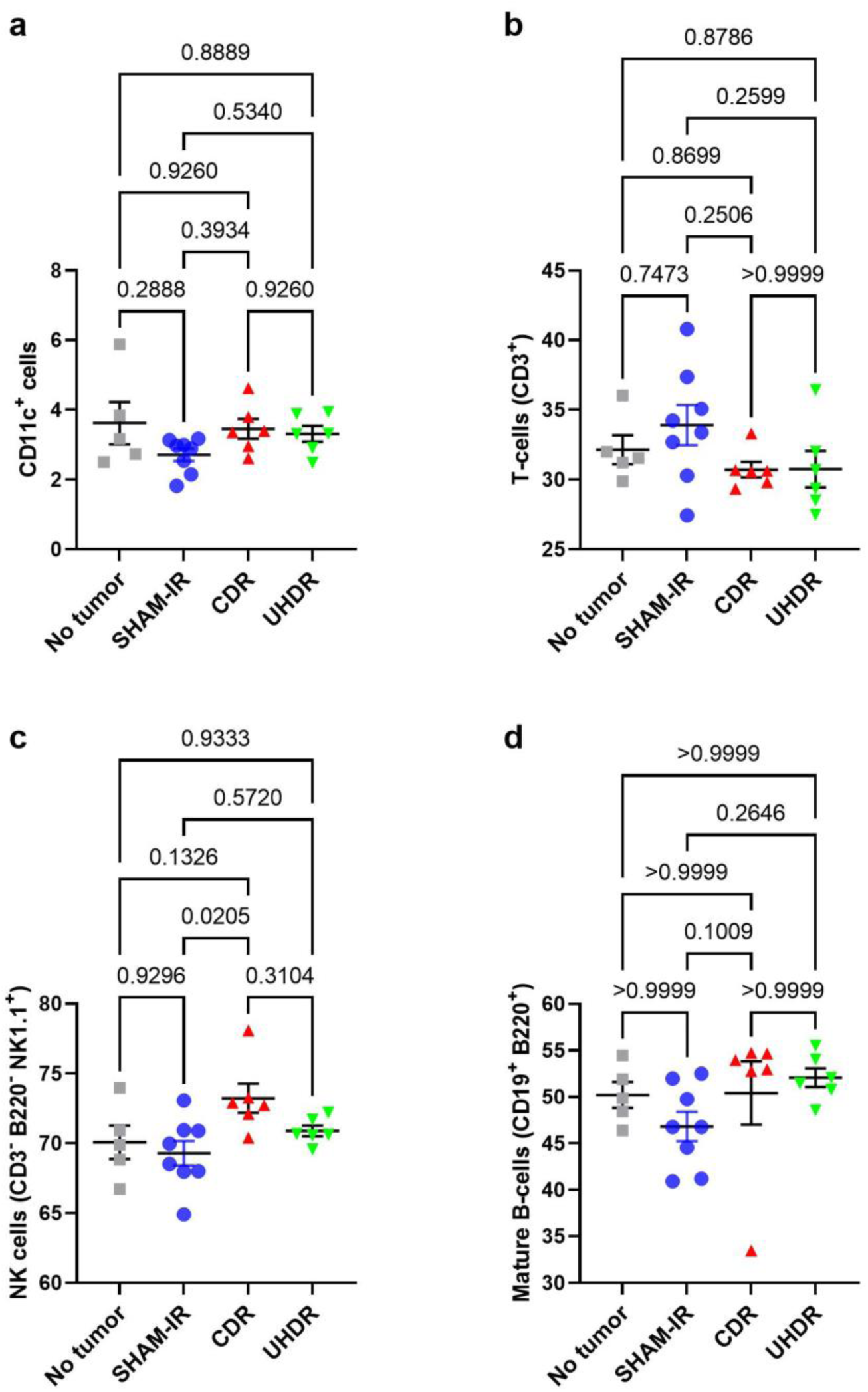
Main splenic immune cell populations 28 days following CDR or UHDR treatment. a-d) Percentages of splenic immune cell populations; No tumor n=5, SHAM-IR n=8, CDR n=6, UHDR n=6. Black bars indicate mean values ± SEM for each treatment group for a) CD11c^+^ cells, b) T-cells, c) NK cells and d) mature B-cells. The p-values (Kruskal-Wallis test) are indicated individually.

**Figure S2.**
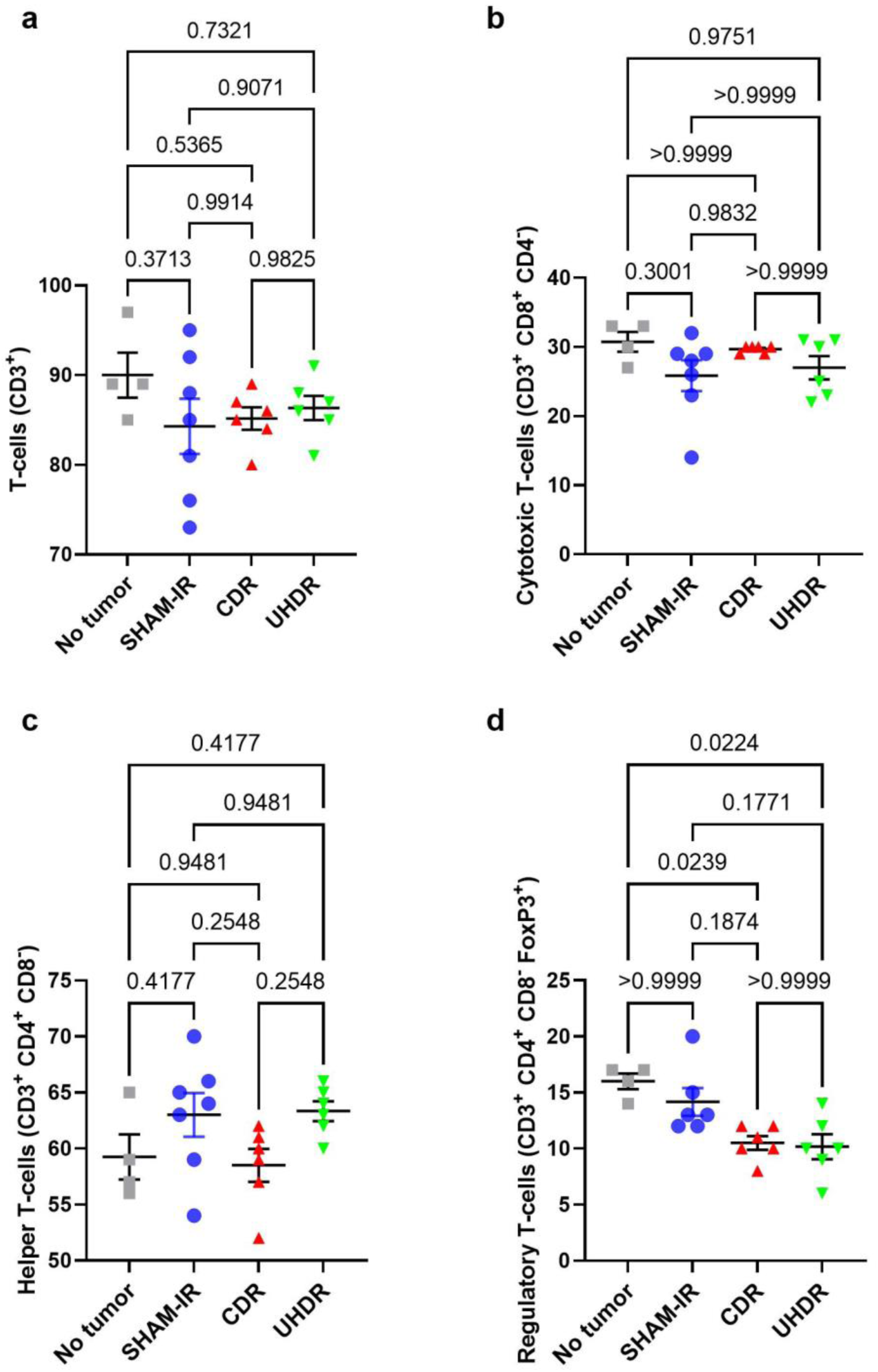
T-cell populations in lymph nodes 28 days following CDR or UHDR treatment. a-d) Percentages of different T-cell populations; No tumor n=5, SHAM-IR n=8, CDR n=6, UHDR n=6. Black bars indicate mean values ± SEM for each treatment group for a) T-cells, b) cytotoxic T-cells, c) helper T-cells and d) regulatory T-cells. The p-values (Kruskal-Wallis test) are indicated individually.

